# Investigating receptor-mediated antibody transcytosis using Blood-Brain Barrier organoid arrays

**DOI:** 10.1101/2021.02.09.430382

**Authors:** Claire Simonneau, Martina Duschmalé, Alina Gavrilov, Nathalie Brandenberg, Sylke Hoehnel, Camilla Ceroni, Evodie Lassalle, Hendrik Knoetgen, Jens Niewoehner, Roberto Villaseñor

## Abstract

**Background:** The pathways that control protein transport across the Blood-Brain Barrier (BBB) remain poorly characterized. Despite great advances in recapitulating the human BBB *in vitro*, current models are not suitable for systematic analysis of the molecular mechanisms of antibody transport. The gaps in our mechanistic understanding of antibody transcytosis hinder new therapeutic delivery strategy development.

**Methods:** We applied a novel bioengineering approach to generate human BBB organoids by the self-assembly of astrocytes, pericytes and brain endothelial cells with unprecedented throughput and reproducibility using micro patterned hydrogels. We designed a semi-automated and scalable imaging assay to measure receptor-mediated transcytosis of antibodies. Finally, we developed a workflow to use CRISPR/Cas9 gene editing in BBB organoid arrays to knock out regulators of endocytosis specifically in brain endothelial cells in order to dissect the molecular mechanisms of receptor-mediated transcytosis.

**Results:** BBB organoid arrays allowed the simultaneous growth of more than 5000 homogenous organoids per individual experiment in a highly reproducible manner. BBB organoid arrays showed low permeability to macromolecules and prevented transport of human non-targeting antibodies. In contrast, a monovalent antibody targeting the human transferrin receptor underwent dose- and time-dependent transcytosis in organoids. Using CRISPR/Cas9 gene editing in BBB organoid arrays, we showed that clathrin, but not caveolin, is required for transferrin receptor-dependent transcytosis.

**Conclusions:** Human BBB organoid arrays are a robust high-throughput platform that can be used to discover new mechanisms of receptor-mediated antibody transcytosis. The implementation of this platform during early stages of drug discovery can accelerate the development of new brain delivery technologies.

## Background

In recent years, developing technology platforms to deliver biologics to the brain parenchyma from the circulation has accelerated. Robust evidence from independent groups shows that targeting receptors present on endothelial cells at the blood-brain barrier (BBB) can lead to enhanced brain uptake of therapeutic proteins in preclinical species [1–5]. The diversity of shuttle formats (e.g. bispecific antibodies [1], single domain antibodies [6–9], Fc-engineered binders [3]) together with different receptor targets in brain endothelial cells (e.g. transferrin receptor [2,10–12], insulin receptor [13], CD98 [14], TMEM30 [15]) expand the design space for successful brain delivery in a combinatorial manner. Despite these rapid advances in protein delivery technologies to the central nervous system (CNS), the understanding of the precise molecular mechanisms of receptor-mediated antibody transport across the BBB is lagging.

Recent studies have shed light on candidate pathways for antibody transcytosis across the BBB. We previously found that intracellular endosome tubules regulate transferrin receptor-based brain shuttle sorting for transcytosis [16]. Sorting tubules are also involved during transcytosis of the low-density lipoprotein receptor–related protein 1 [17] and a specific serotype of adeno-associated virus (AAV9) [18]. In contrast, FC5 transcytosis depends on multivesicular body transport [15]. These results highlight that receptor-mediated transcytosis may occur via a variety of independent pathways. Identifying the molecular regulators of these transport pathways requires functional genomic tools to evaluate the effect of individual genes on transcytosis, as was recently shown for polarized epithelial cells [19]. However, most work on the regulation of transcytosis at the BBB is still performed in rodents [20–22]. While these landmark efforts identified novel regulators for specific pathways, such as Mfsd2a [20] and ALPL [23], *in vivo* studies do not enable sufficiently high throughput and are not optimal to perform unbiased system-level analysis of transport mechanisms.

Independent groups have recently established a novel organoid-based model of the BBB formed by the self-organization of human brain endothelial cells, pericytes and astrocytes [24–26]. BBB organoids allow direct cell-cell interactions in the absence of artificial membranes or substrates and, importantly, recapitulate key cellular and molecular properties of the BBB, including: 1) tight junction formation, 2) efflux pump expression and activity and 3) receptor-mediated transport of peptides [26]. Although this system is a major advance in the field of *in vitro* BBB modelling, organoid production, processing and analysis still require extensive manual work [27]. Thus, the complexity of the protocol and its relative low-throughput hinder its widespread adoption for screening purposes.

Here, we used a recently developed technology based on hydrogel micro-scaffolds [28] to generate patterned BBB organoid arrays, which allowed the formation of more than 5000 viable organoids with reproducible sizes in a 96-well plate format. In agreement with previously described human *in vitro* models, BBB organoid arrays restricted the passage of large molecules to the organoid core but recapitulated receptor-mediated transcytosis of a transferrin receptor-targeting antibody shuttle. Finally, we showed that BBB organoids can be combined with CRISPR/Cas9 gene editing to knock out specific genes in brain endothelial cells to evaluate their role in receptor-mediated transcytosis. Together, our data show that BBB organoid arrays are a high-throughput platform to discover new mechanisms of antibody transport across the BBB.

## Methods

### Fabrication of hydrogel-based U-bottom microwell arrays

U-bottom microwell hydrogel-based arrays (Gri3D, SUN bioscience) were fabricated and conditioned as previously described [28]. For the generation of high-throughput BBB organoids, microcavities of 600 μm in diameter and 720 μm in depth imprinted in polyethylene glycol (PEG) hydrogels (Elastic modulus: G’= 12.5 kPa) were typically used.

### Culture conditions

Primary human astrocytes (HA, ScienCell Research Laboratories) were grown in Astrocyte Growth Medium (AGM) composed of Astrocyte Medium (AM, ScienCell Research Laboratories) supplemented with 2% FBS, 1% Astrocytes Growth Supplement (AGS, ScienCell Research Laboratories) and 1% penicillin/streptomycin. Human brain microvascular pericytes (HBVP, ScienCell Research Laboratories) were cultured in Pericyte Growth Medium (PGM) composed of Pericytes Medium (PM, ScienCell Research Laboratories) supplemented with 2% FBS, 1% Pericytes Growth Supplement (PGS, ScienCell Research Laboratories) and 1% penicillin/streptomycin. Human cerebral microvascular endothelial cells (hCMEC/D3, Sigma-Aldrich) were maintained in culture using Endothelial Basal Medium (EBM-2, Lonza) supplemented with hydrocortisone, GA-1000, 5% FBS, hEGF, VEGF, hFGF-B, R3-IGF-1, ascorbic acid and heparin (EGM-2 SingleQuots Supplements, Lonza). Cells were grown in T-75 flasks, coated with 2 μg/cm^2^ of poly-L-lysine for HBVP and HA. The growth medium was changed every two days. For experimental use, cells were grown until 90% confluence before subculturing; HA and HBVP were maintained between passages p1 and 4 and hCMEC/D3 cells were used for 10 passages. For the generation of BBB organoids, EGM-2 medium without VEGF and reduced FBS (2%) was used, and named hereinafter Organoid Medium (OM).

### BBB organoid generation

HA, HBVP and hCMEC/D3 were detached by 0.05% trypsin/EDTA (ThermoFisher Scientific) and resuspended in warm OM. The concentration of each cell type was determined using the Countess™ automated cell counter (Invitrogen). The cells were resuspended at the appropriate concentration to target a thousand cells per microwell in a seeding volume of 60 μL per well (a total of 3000 cells per microwell in a 1:1:1 ratio). The cells were grown in a humidified incubator at 37 °C with 5% CO_2_ for 48 hours to allow self-assembly of the multicellular organoids.

### Imaging and size analysis of BBB organoids

The assembled multicellular BBB organoids were imaged for size analysis after 48 hours on a Nikon Eclipse Ti-E fully motorized microscope (Nikon) by automatically stitching 4×4 field of view images using a CFI Plan Fluor DL 4X N.A. 0.13, W.D. 16.4mm, PH-L air objective. All bright field images were analyzed and quantified using a custom-made Python script to estimate the diameter and size homogeneity of individual organoids.

### Immunofluorescence staining of BBB organoids

Multicellular BBB organoid arrays were fixed in 4% paraformaldehyde for 45 minutes at room temperature. Samples were washed thoroughly with PBS and were permeabilized and blocked with 0.6% Triton-X + 10% donkey serum in PBS for 1 hour at room temperature. Organoids were then transferred to Eppendorf protein LoBind tubes for staining. All antibodies used in this study are reported in Table 1. Primary antibodies were then added in dilution buffer (0.1% Triton-X + 10% Donkey serum in PBS) and incubated overnight at 4 °C. Organoids were washed thoroughly in washing buffer (0.1% Triton-X in PBS) and incubated overnight at 4 °C with the respective species-specific fluorescently-labelled antibodies and DAPI (1 μg/mL, Sigma-Aldrich). Finally, the samples were washed again in the washing buffer and transferred into a 96 glass-bottom imaging plate containing 50 μL of Fluoromount per well. Samples for qualitative evaluation of marker localization were imaged using a Zeiss LSM700 inverted confocal microscope (Zeiss) using an EC Plan-NeoFluor 10X N.A. 0.3, W.D. 5.2mm, PH 1 air and Plan-Apochromat 20X N.A. 0.8, W.D. 0.55mm, PH 2 air objective.

**Table 1.**
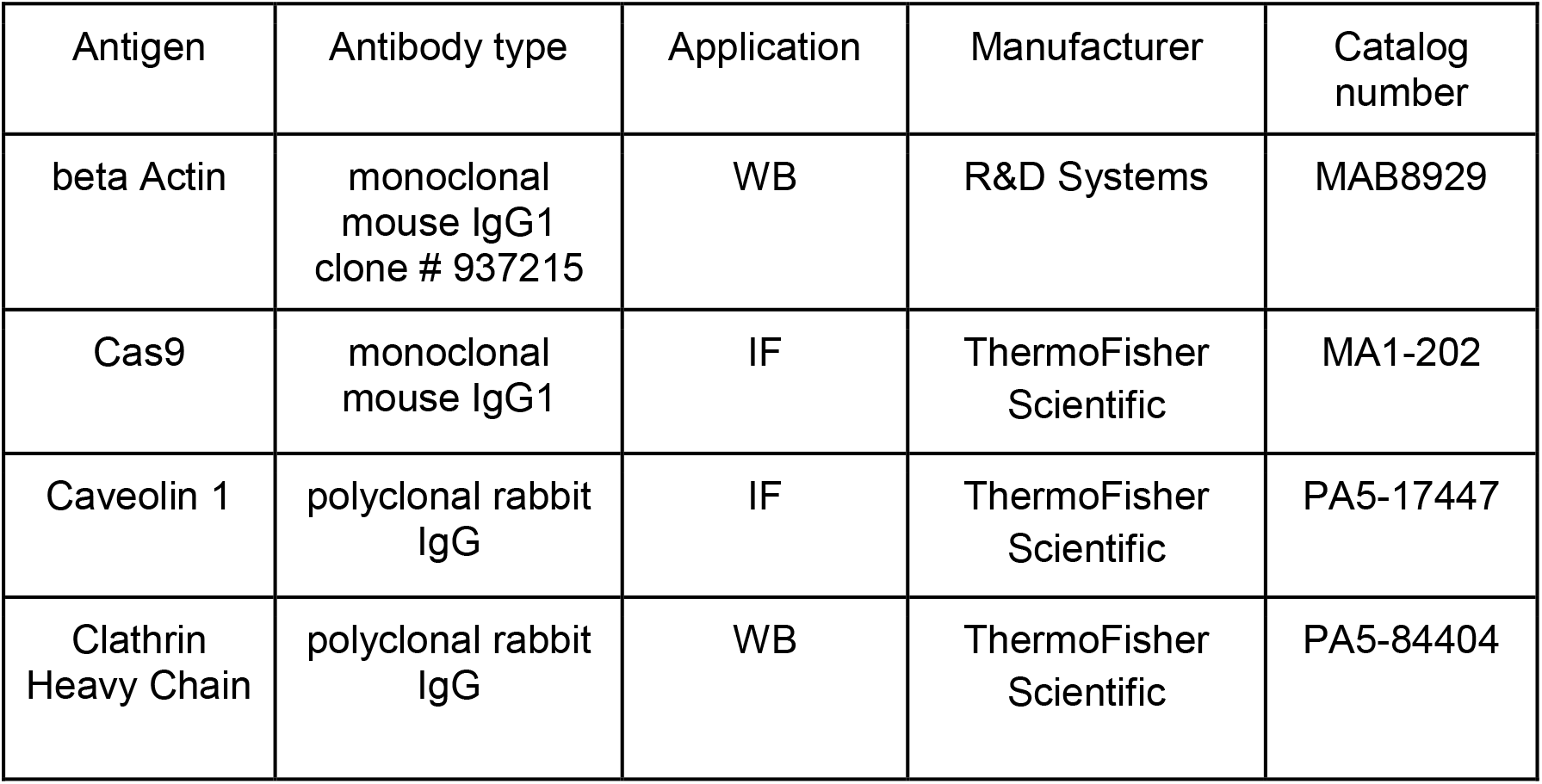
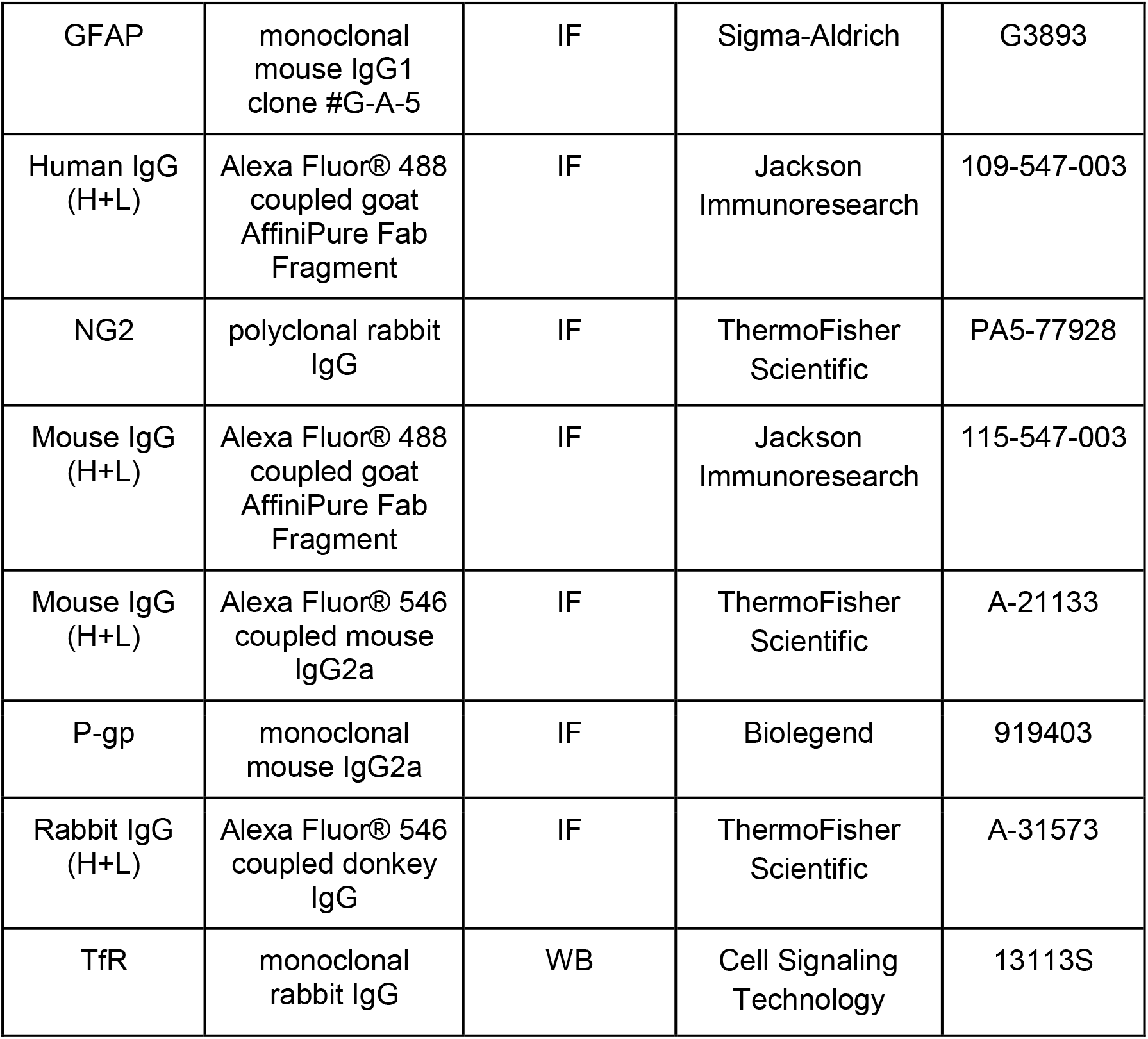
List of antibodies.

### Antibody list

### Live/dead assay

After 48 hours of assembly, BBB organoids were labelled with calcein-AM and ethidium homodimer-1 according to the manufacturer’s protocol (Live/DEAD Viability/Cytotoxicity Kit for mammalian cells, ThermoFisher Scientific) and were imaged using a a Nikon Eclipse Ti-E fully motorized microscope (Nikon).

### Dextran permeability assay

Multicellular BBB organoid arrays were formed for 48 hours and were incubated with FITC-Dextran of different molecular weight (4, 40 and 70 kDa, Sigma-Aldrich) in OM for 4 h at 37 °C with 5% CO_2_. Organoids were washed 3 times with PBS, and fixed in 4% paraformaldehyde for 30 minutes at room temperature. The organoids were transferred to Eppendorf protein LoBind tubes and stained with DAPI (1 μg/mL, Sigma-Aldrich). Finally, organoids were washed with washing buffer (0.1% Triton-X in PBS), transferred to cover glasses, and mounted with Fluoromount (Electron Microscopy Science). Organoids were imaged using a Leica SP5 confocal microscope using an HCX PL APO CS 40X/1.3 oil objective (Leica). The organoid core was defined as the volume starting at 25 μm depth from the top of the organoid. Z-stacks covering a total depth of 3.5 μm were acquired for quantification. Quantification of organoid permeability to FITC-dextran was performed using a custom-made automated Fiji [29] script that segments individual organoids and measures the mean fluorescence intensity of the maximum intensity projection within 75% of the cross-section area at the spheroid core.

### Transcytosis assay with BBB organoid arrays

After 48 hours of assembly, BBB organoid arrays were incubated with custom-made humanized monovalent antibodies targeting either the human or the mouse transferrin receptor [2] and a non-targeting human IgG as a control in OM for 4 hours at 37 °C with 5% CO_2_. BBB organoids were washed thoroughly 6 times for 5 minutes with pre-warmed OM in the incubator. Samples were then fixed, processed for immunostaining and imaged for quantitative analysis as described above.

### Gene editing in BBB organoids

Ready-to-use lentiviral particles were purchased from Sigma-Aldrich. hCMEC/D3 cells were transduced first by lentiviral particles expressing Cas9 and blue fluorescent protein (BFP) as a transduction marker (LVCAS9BST, Sigma-Aldrich). Cas9-expressing cells were then selected with blasticidin antibiotic at 50 μg/mL for 6 days (Life technologies). Cas9-expressing hCMEC/D3 cells were each transduced separately by lentiviral particles expressing sgRNAs and selected by puromycin resistance at 10 μg/mL for 6 days (Life technologies). sgRNA sequences are listed in Table 2.

**Table 2.**
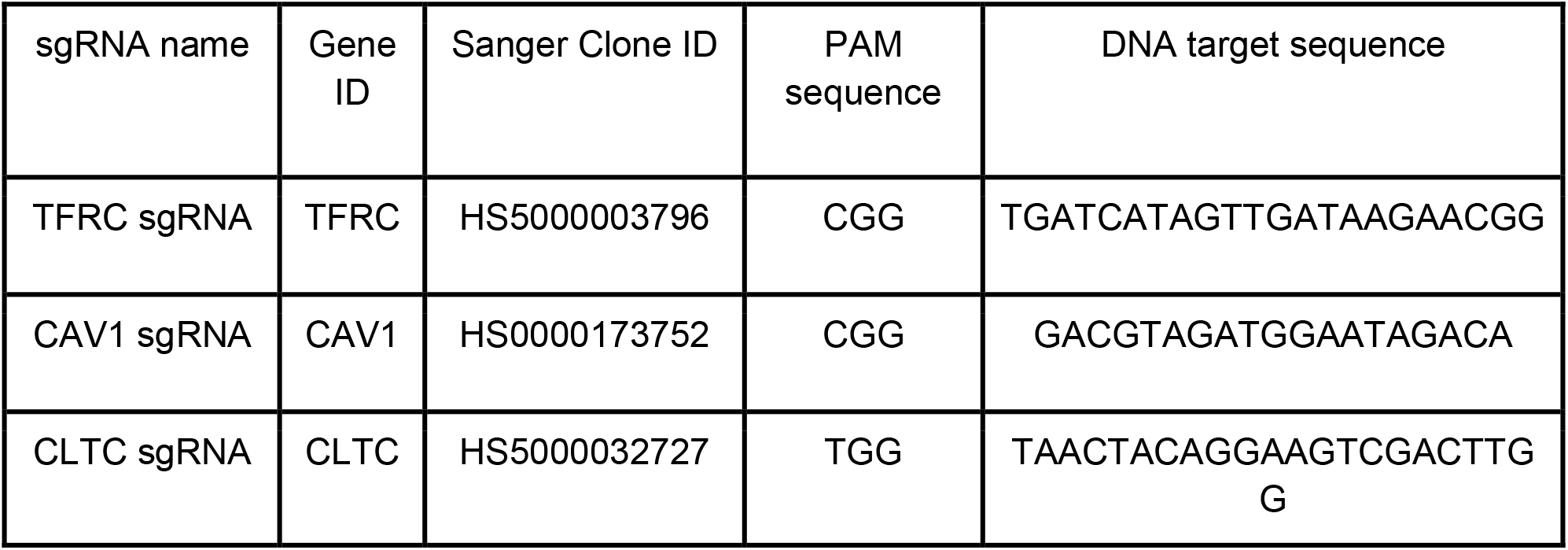
List of sgRNAs.

To generate monoclonal KO cell populations, Cas9-gRNA-expressing hCMEC/D3 were sorted (FACSAria II, BD Biosciences) using BFP into single clones and subsequently expanded.

### Western blot analysis

Whole cell lysates from hCMEC/D3 were prepared in RIPA lysis buffer (ThermoFisher Scientific) supplemented with cOmplet Protease Inhibitor Cocktail (Roche). Lysates were clarified by centrifugation (20000 g, 10 minutes, 4 °C) and protein concentrations were determined using the BCA protein assay kit (ThermoFisher Scientific). The same amount of protein (1.2 μg) was separated and analyzed for protein level using the 12-230 kDa Wes Separation Module with the Wes system (Protein Simple). All primary antibodies used are reported in Table 1. Secondary antibodies were used following the manufacturer recommendations (Protein Simple).

### Transferrin uptake

hCMEC/D3 cells were grown on pre-coated 24-well glass bottom plates (Mattek) with collagen I (250 μg/mL, Sigma-Aldrich) for 48 hours. To examine transferrin uptake, cells were incubated at 37 °C with 5% CO_2_ with Alexa Fluor 488–conjugated transferrin (25 μg/mL, ThermoFisher Scientific) for 30 minutes. Cells were quickly washed with PBS, fixed in 4% paraformaldehyde for 20 minutes at room temperature and permeabilized and blocked in 0.1% saponin (Sigma-Aldrich) + 4% gelatin from cold water fish skin (Sigma-Aldrich) in PBS for 10 minutes at room temperature. Cells were then incubated with phalloidin atto-565 (200 pM, Sigma-Aldrich) and DAPI (1 μg/mL, Sigma-Aldrich) for 30 minutes at room temperature and washed 3 times with PBS. Cells were imaged with a DMi8S Leica widefield microscope using a HC PL APO 100X/1.47 oil objective. Quantification of total vesicle fluorescent intensity was made using MotionTracking as previously described [30,31].

### Statistics

All values from quantifications are shown as box plots with median and interquartile ranges with lines representing the 5th and 95th percentiles. Statistical comparisons between multiple groups were performed by one-way analysis of variance (ANOVA), followed by Dunnett’s post hoc test. Values were considered to be significantly different when p < 0.05. Statistical analysis and plotting of data were performed with GraphPad Prism Software version 8.

## Results

### Assembly of BBB organoids in hydrogel microwell arrays

To scale up the generation of BBB organoids, we used micropatterned hydrogel Gri3D plates, which allow the generation of thousands of organoids per experiment (Figure 1a) [28]. Specifically, we co-cultured primary human astrocytes (HA) and human brain vascular pericytes (HBVP) with immortalized human brain cerebral microvascular endothelial cells (hCMEC/D3) in U-bottom microwell hydrogel-based inserts placed on custom-made 96-well plates. All three cell types were mixed in a 1:1:1 ratio and seeded on the hydrogel microcavity arrays to target an average of 3000 cells per microcavity. After 48 hours, cells formed compact homogeneous organoids throughout the plate at predefined locations and on the same focal plane (Figure 1a). To assess the reproducibility of organoid arrays between experiments, we analyzed the diameter of BBB organoids generated in GRi3D microwell arrays and compared it to the previously reported method of organoid formation on agarose substrate [27]. In both cases the average diameter was comparable (220 −250 um) and in agreement with previously reported data [24,26] (Figure 1b). The diameter of BBB organoids in arrays showed higher intra-experiment variability (SD between 23 and 27) compared to agarose (SD between 6 and 22). However, organoid diameter in Gri3D plates was highly reproducible across 3 independent experiments, whereas in agarose the diameter varied up to 40% (Figure 1b). Importantly, the average BBB organoid size was not affected by the specific Gri3D plate format, as we observed the same results using hydrogels in 24-well plates with the same microwell diameter (Figure 1b). This demonstrates that BBB organoid assembly can be upscaled using larger arrays without affecting its reproducibility.

**Figure 1.**
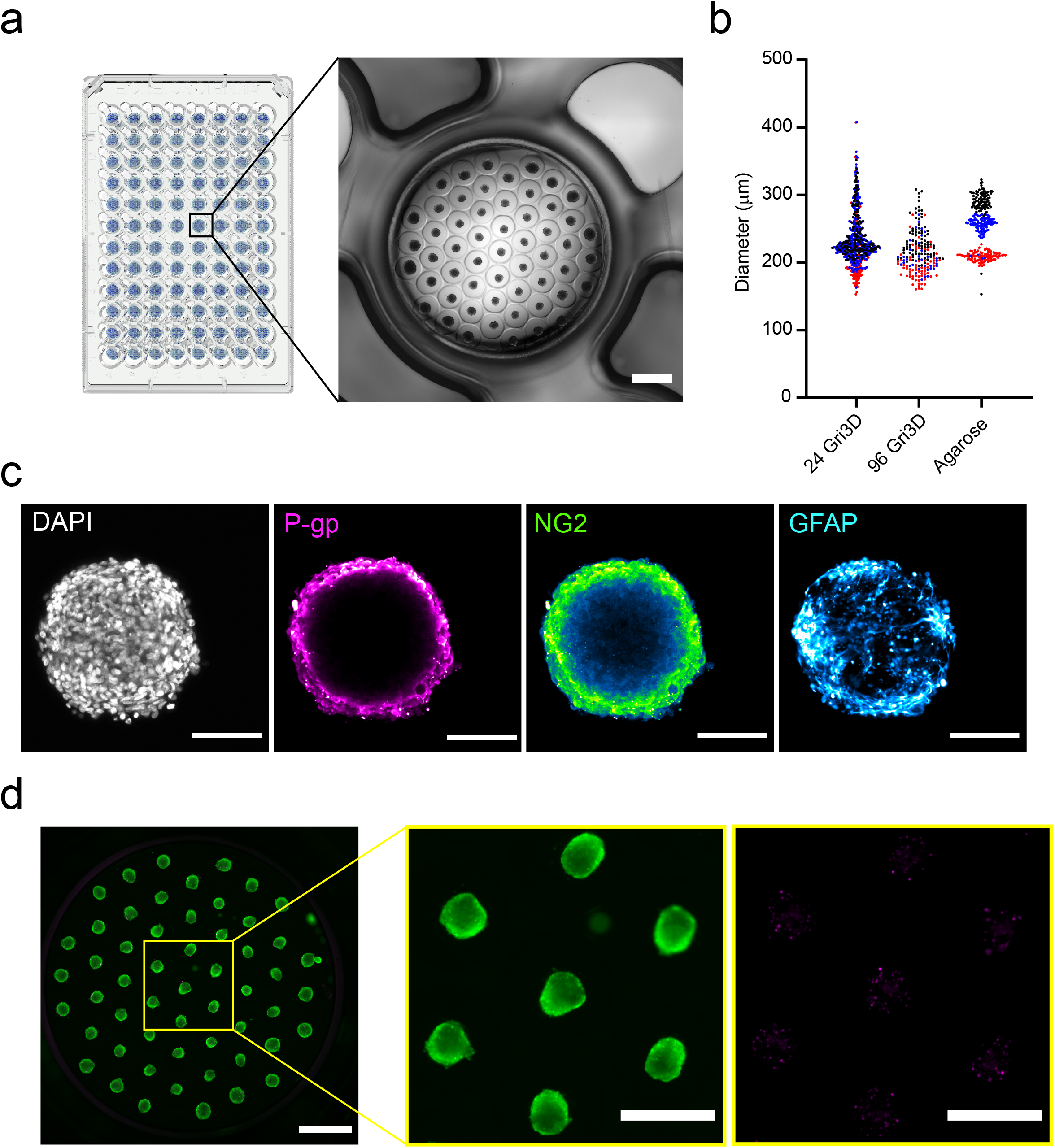
High-throughput blood-brain barrier organoid formation with microwell hydrogel arrays. **a,** Schematic of microwell plates in 96-well plate format and representative phase contrast image of a GRi3D organoid array within a single well. Scale bar, 1 mm. **b,** Quantification of blood-brain barrier organoid diameter in GRi3D microwell arrays within 24- or 96-well plate compared to organoids grown on agarose substrate. Each dot represents a single organoid, while different colors represent independent experiments. **c,** Representative confocal images of a blood-brain barrier organoid labelled with cell type-specific markers showing specific layering and organization of brain endothelial cells (P-gp, magenta), pericytes (NG2, green) and astrocytes (GFAP, cyan) when grown in GRi3D arrays. Nuclei are labelled with DAPI (grey). Scale bars, 100 μm. **d,** Representative fluorescent image of a live/dead staining showing the viability of blood-brain barrier organoids grown on GRi3D arrays. Calcein AM in green labels live cells whereas ethidium homodimer-1 in magenta labels dead cells. Scale bar, 1 mm. Images on the right show a higher magnification of the boxed area. Scale bar 500 μm.

Next, we confirmed self-assembly and patterning of individual cell types into well-organized neurovascular structures by performing immunofluorescence stainings on compact BBB organoids grown in microwell arrays. We used cell-specific markers to identify each cell type in the model; endothelial cells were tested for P-gp expression, pericytes for neural/glial antigen 2 expression (NG2) and astrocytes for glial fibrillary expression (GFAP). After 48 hours of culture, we found that the brain endothelial cells (BECs) formed a distinct monolayer at the surface of the organoids, in close contact with a layer of pericytes, which separate the BECs from the astrocytes located in the core of the organoids (Figure 1c). This specific multicellular self-organization recapitulates the native cell-cell interactions within the BBB *in vivo* and is consistent with previous BBB organoid protocols, which used high-resolution imaging to show expression of BBB markers [24–27]. Finally, we examined whether the high number of BBB organoids within the limited media volume in a single well would affect cell viability after 48 hours of assembly. We labeled the BBB organoids with calcein-AM and ethidium homodimer-1 to detect viable and dead cells. We found that, under these conditions, the vast majority of cells in BBB organoids were viable and only a few dead cells were scattered across the organoid (Figure 1d). Therefore, micropatterned hydrogel arrays offer a suitable alternative for reproducible growth of self-assembled and viable BBB organoids at a large scale.

### Antibody receptor-mediated transcytosis in BBB organoid arrays

Next, we designed a standardized imaging workflow to estimate large molecule transport across organoids. We used P-gp expression to identify and distinguish the organoid surface from the organoid core (Figure 2a). We reduced acquisition time by capturing a limited confocal volume (3.5 um in z) within the core of each individual organoid. These imaging data sets were analyzed with a Fiji script that automatically segments BBB organoids and quantifies fluorescence intensity at the center of the organoid cross section (Figure 2b). To test this accelerated imaging workflow and automated analysis, we first assessed the barrier function of BBB organoid arrays by incubating organoids with fluorescently labeled dextran of different molecular weights (4, 40 and 70 kDa). In agreement with previous findings, we could not detect FITC-dextran within the organoid core, and the residual fluorescent signal was comparable with that of non-treated BBB organoids (Figure 2c). This qualitative observation was further supported by the automatic quantification of a larger population of BBB organoids across independent experiments with the image analysis script (Figure 2d). This result shows that BBB organoid arrays maintain the low paracellular permeability to macromolecules reported in other *in vitro* models [24–27,32].

**Figure 2.**
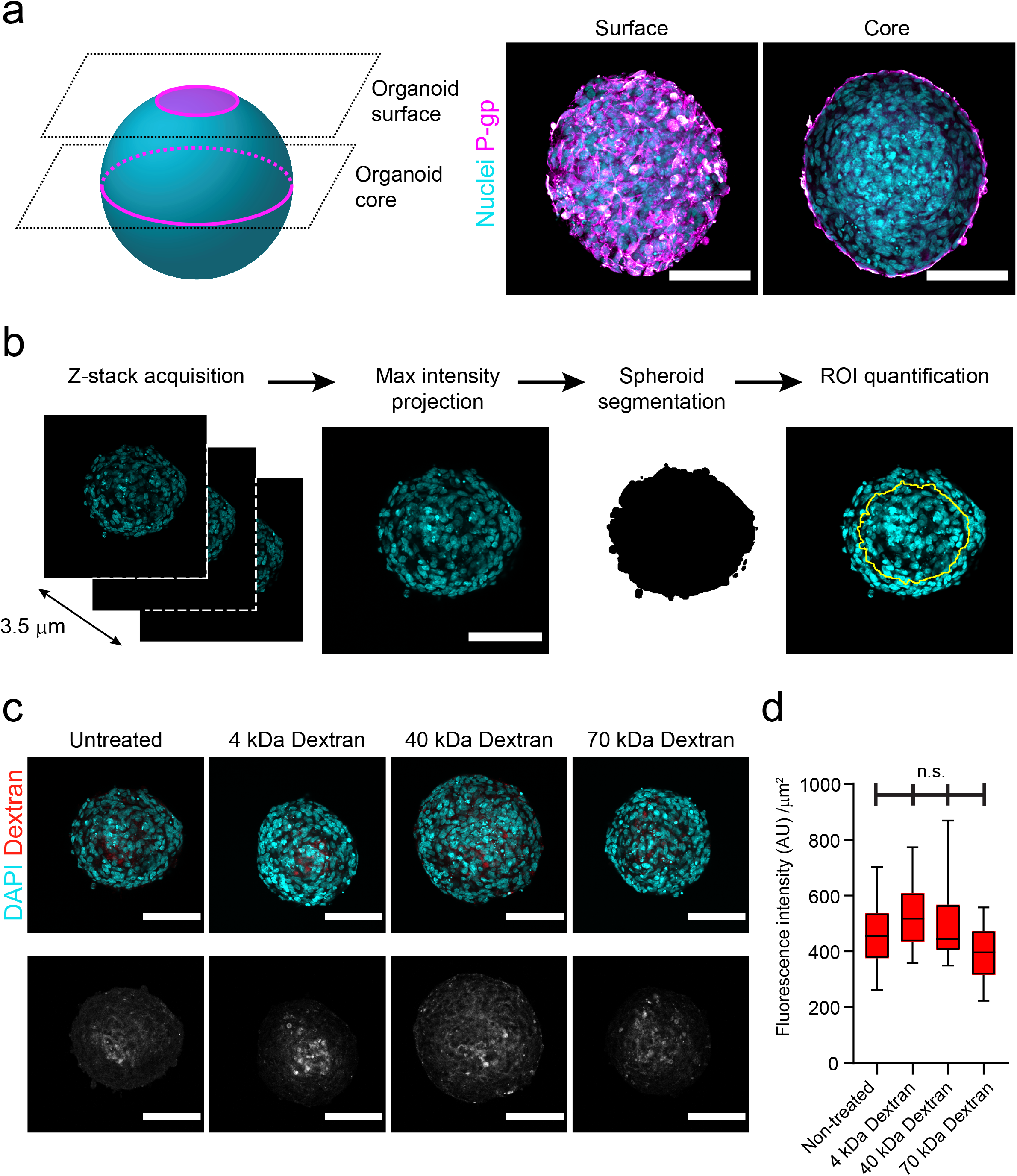
Standardized imaging workflow to assess transport into blood-brain barrier organoids. **a,** Scheme showing the relative position of imaging planes for experimental acquisition (left) and representative confocal images (right) acquired at the organoid surface and core positions. Brain endothelial cells are labelled with an anti-P-gp antibody (magenta) and are homogeneously distributed at the surface and form a continuous monolayer at the periphery of the organoid core. Nuclei are labelled with DAPI (cyan). Scale bar, 100 μm. **b,** Scheme representing the Z-stack acquisition and analysis process (maximum intensity projection, segmentation and ROI definition) to quantify fluorescent molecules within the organoid core. In all images, nuclei are labelled with DAPI (cyan). Scale bar, 100 μm. **c,** Representative confocal images of blood-brain barrier organoids incubated with different molecular weight Dextrans for 4 hours. The upper images show an overlay of Dextran (red) and nuclei labelled with DAPI (cyan). The intensity of the lower images was scaled to visualize the background intensity in the Dextran channel (grey). Scale bars, 100 μm. **d,** Quantification of Dextran fluorescence intensity within blood-brain barrier organoids. Graph shows boxplots with interquartile ranges and median. Lines show the 5th and 95th percentiles. Differences between treatments were not statistically significant (p = 0.51) as evaluated by one-way ANOVA of 50 organoids per condition in n = 4 independent experiments.

Next, we evaluated the BBB organoid arrays as a screening platform for receptor-mediated transcytosis with this imaging workflow. We used a monovalent antibody against the extracellular domain of the human transferrin receptor (TfR) (human Brain Shuttle) [2]. We incubated BBB organoid arrays with 100 nM of non-labeled antibody and detected it after 4 hours with fluorescently-labeled anti human IgG-specific Fab fragments. As negative controls, we used a non-targeting human IgG and a monovalent antibody against the mouse TfR that does not cross-react with the human receptor (mouse Brain Shuttle). Under these conditions, we did not detect either the non-targeting IgG or the mouse Brain Shuttle within BBB organoids. In contrast we detected substantial accumulation of human Brain Shuttle within the organoid core (Figure 3a, b). As evidenced by quantification of more than 50 individual BBB organoids per condition, transport of human Brain Shuttle into BBB organoids occurred in a dose- and time-dependent manner (Figure 3c, d). Overall, these data show that BBB organoid arrays combined with this optimized imaging and analysis workflow are a suitable platform for identifying and characterizing antibodies that undergo receptor-mediated transcytosis across the human BBB.

**Figure 3.**
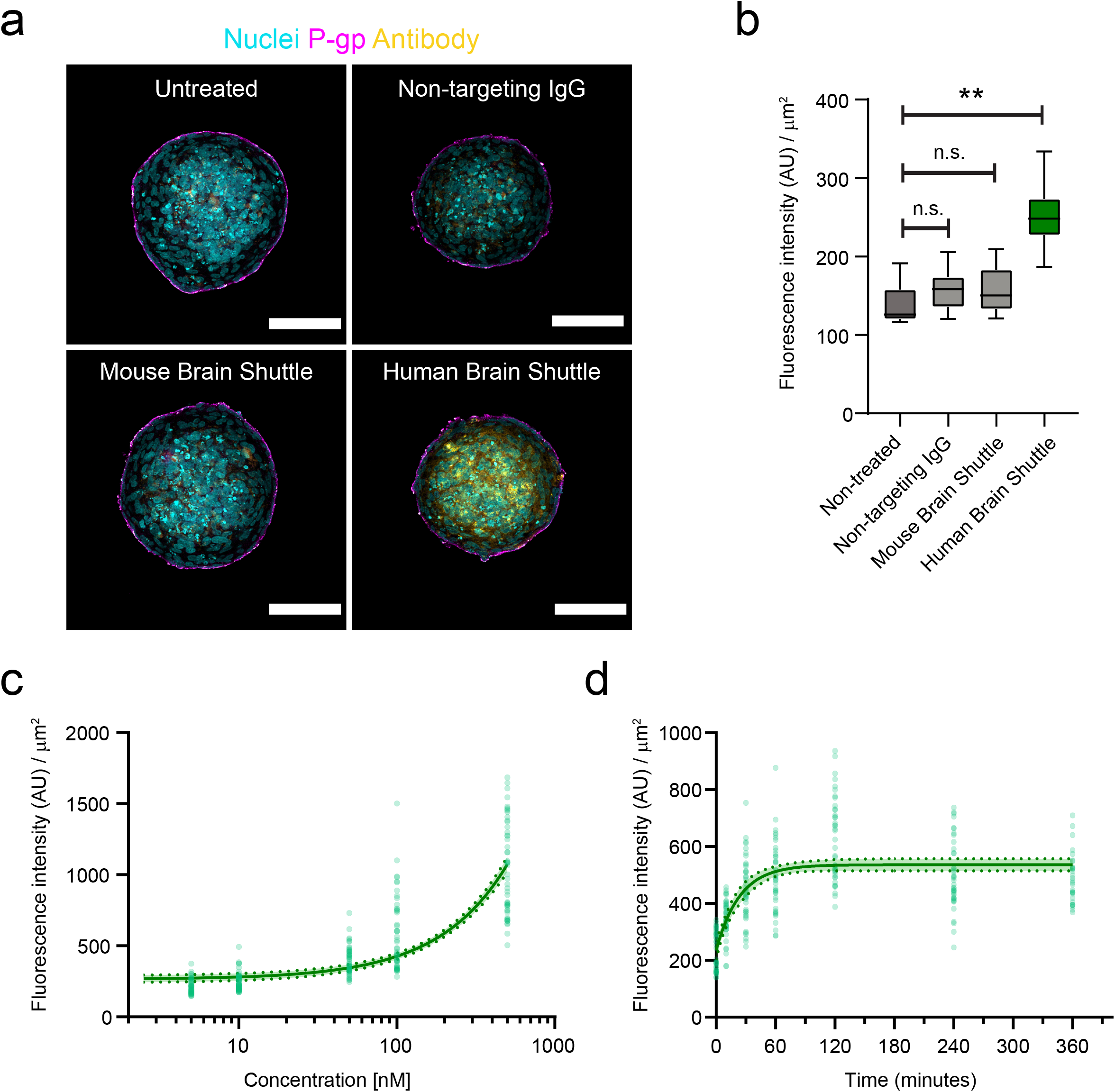
Receptor-mediated transcytosis of a human Brain Shuttle in blood-brain barrier organoids. **a,** Representative confocal images of blood-brain barrier organoids after incubation with different antibody constructs for 4 hours. Brain endothelial cells are labelled with an anti-P-gp antibody (magenta), antibody constructs are labelled with anti-human IgG Fab fragments (yellow) and nuclei with DAPI (grey). Scale bars, 100 μm. **b,** Quantification of antibody construct transcytosis into blood-brain barrier organoids. Graph shows boxplots with interquartile ranges and median. Lines show the 5th and 95th percentiles. Differences between non-treated organoids and non-targeting IgG or mouse brain shuttle were not statistically significant (p > 0.6) whereas differences between human brain shuttle and non-treated control were statistically significant (**, p = 0.002) as evaluated by one-way ANOVA followed by Dunnett’s test for multiple comparisons of 60 organoids per condition in n = 3 independent experiments. **c,** Dose-response of human brain shuttle transcytosis into the core of blood-brain barrier organoids. **d,** Time course of human brain shuttle transcytosis into the core of blood-brain barrier organoids. In **c** and **d** points represent individual organoids dosed at different concentrations. The solid lines show the best linear (**c**) or nonlinear (**d**) fit of the experimental data. The dotted lines and colored area show the 95% confidence interval of the curve estimation.

### Dissecting the molecular regulation of transcytosis with BBB organoid arrays

As BBB organoid arrays enable robust and sensitive transport assays, we further investigated the mechanisms of receptor-mediated transcytosis with CRISPR-based gene editing. In order to interrogate the role of specific genes on transcytosis across the BBB, we first generated a human brain endothelial cell line (hCMEC/D3) stably expressing Cas9. Importantly, this Cas9-expressing cell line maintained its capacity to self-organize at the surface of BBB organoids (Figure 4a). To validate the use of CRISPR-based gene editing on BBB organoid arrays and as a proof-of-concept, we selected genes with well-characterized functions in transferrin transport and/or endocytosis: TfR, clathrin heavy chain and caveolin-1 [33–35]. We used a non-targeting scrambled sgRNA as a negative control. To functionally characterize these cell lines we analyzed protein expression and quantified transferrin uptake, a well-established assay to evaluate perturbations to endocytosis [30]. Western blot analysis showed that target protein expression was substantially reduced in all three cell lines compared to the parental and control cells (Figure 4b). In the internalization assay, we observed accumulation of transferrin at the perinuclear region in control cells. On the other hand, transferrin internalization was abrogated in both TfR and clathrin heavy chain knockout cells (Figure 4c, d), in agreement with the essential role of both genes for transferrin uptake [35, 36]. It should be noted that we observed across multiple experiments a fraction of TfR knockout cells that still internalized transferrin, likely reflecting either a small heterozygous population or aggregate, non-functional transferrin taken up by non-specific endocytosis. In contrast, the extent of transferrin internalization and its intracellular localization was not changed in caveolin-1 knockout cells compared to the control (Figure 4c, d). This result confirms that the primary pathway for transferrin internalization in brain endothelial cells is via clathrin-mediated endocytosis [16].

**Figure 4.**
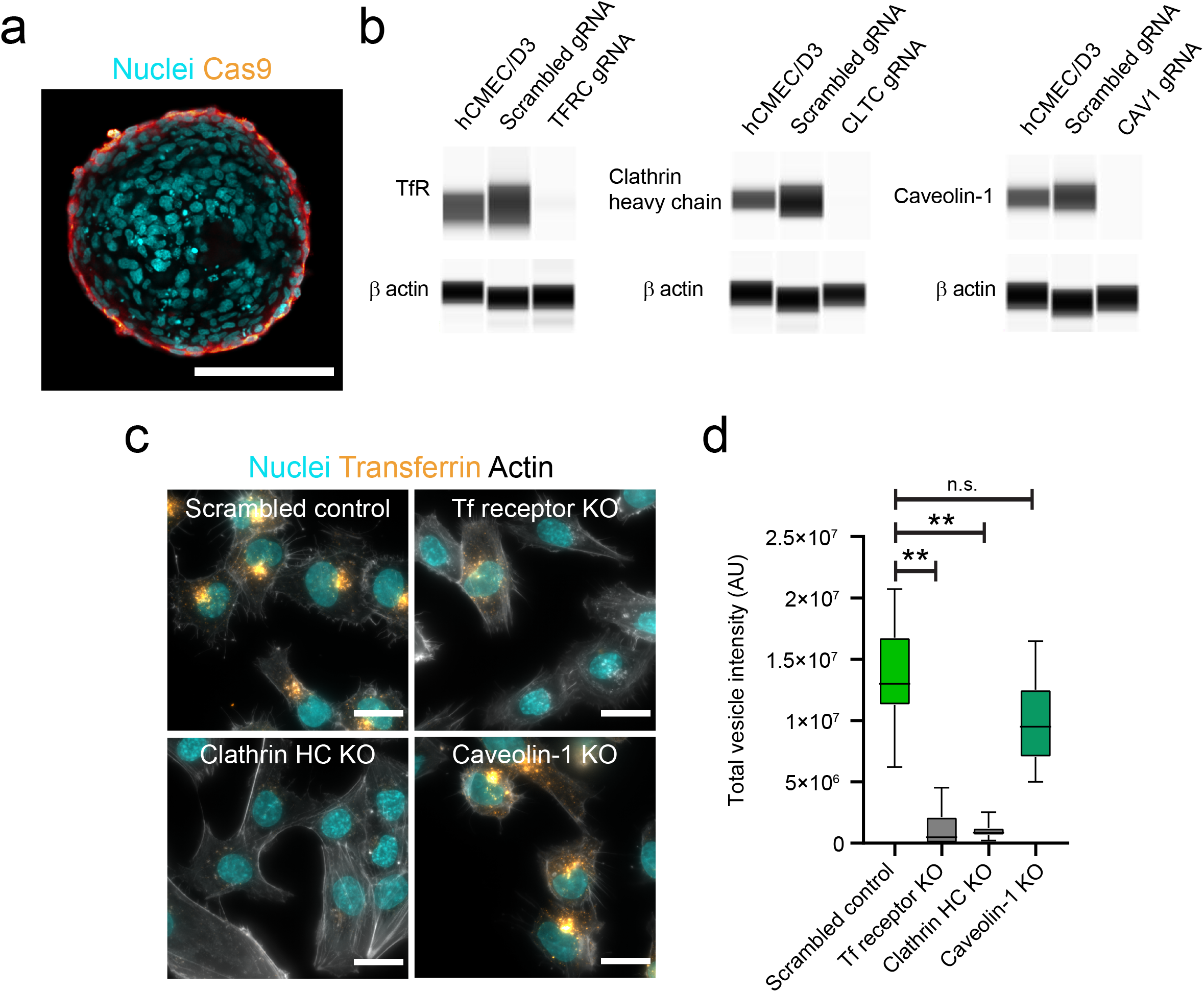
Establishment and characterization of Cas9 brain endothelial cell lines. **a,** Representative confocal image acquired at the core of a blood-barrier organoid assembled with brain endothelial cells expressing Cas9 (orange). Cas9 brain endothelial cells localize only at the periphery of blood-brain barrier organoids. Nuclei labelled with DAPI (cyan). Scale bar, 100 μm. **b,** Representative Western blot images showing the expression of Transferrin receptor, clathrin heavy chain or caveolin-1 in hCMEC/D3 brain endothelial cells expressing Cas9 and transduced with either scrambled gRNA or gRNA against the target gene. β actin expression is shown as a reference control gene. **c,** Representative fluorescent images of hCMEC/D3 brain endothelial Cas9 or knockout cells after incubation with fluorescently labelled transferrin (yellow) for 30 minutes. Actin is labelled with phalloidin (grey) to visualize cell contours and nuclei are labelled with DAPI (cyan). Scale bars, 20 μm. **d,** Quantification of transferrin internalization in hCMEC/D3 Cas9 or knockout cells. Graph shows boxplots with interquartile ranges and median. Lines show the 5th and 95th percentiles. Differences between the scrambled control and transferrin receptor or clathrin heavy-chain knockout cells were statistically significant (**, p = 0.018) whereas the difference between the scrambled control and caveolin-1 knockout cells was not statistically significant (p = 0.418). Comparisons were evaluated by one-way ANOVA followed by Dunnett’s test for multiple comparisons of ~ 400 single cells per condition in n = 2 independent experiments.

We next used each of these knockout cell lines to form BBB organoid arrays and assessed their impact on antibody receptor-mediated transcytosis. Similar to parental hCMEC/D3 or Cas9 cells, all knockout cell lines maintained their capacity to assemble at the surface of BBB organoids (Figure 5a). Furthermore, we could not detect non-targeting IgG within the core of BBB organoids assembled with TfR, clathrin heavy chain or caveolin-1 knockout cells (Figure 5b, c), strongly suggesting that these genes do not affect BBB paracellular permeability. Finally, we incubated knockout organoids with human Brain Shuttle to evaluate receptor-mediated transcytosis. In control organoids, human Brain Shuttle accumulated within the organoid core, whereas in TfR and clathrin heavy-chain knockout organoids the fluorescent signal was substantially reduced and comparable to background levels (Figure 5d, e). Caveolin-1 knockout had no major impact on the transport of human Brain Shuttle into the organoid core (Figure 5d, e). These data show that transcytosis of the human Brain Shuttle across the BBB is TfR-specific and requires clathrin but occurs independently of caveolin-1. Overall, our results demonstrate that BBB organoid arrays can be combined with CRISPR gene editing to interrogate the molecular mechanisms of transcytosis across the human BBB.

**Figure 5.**
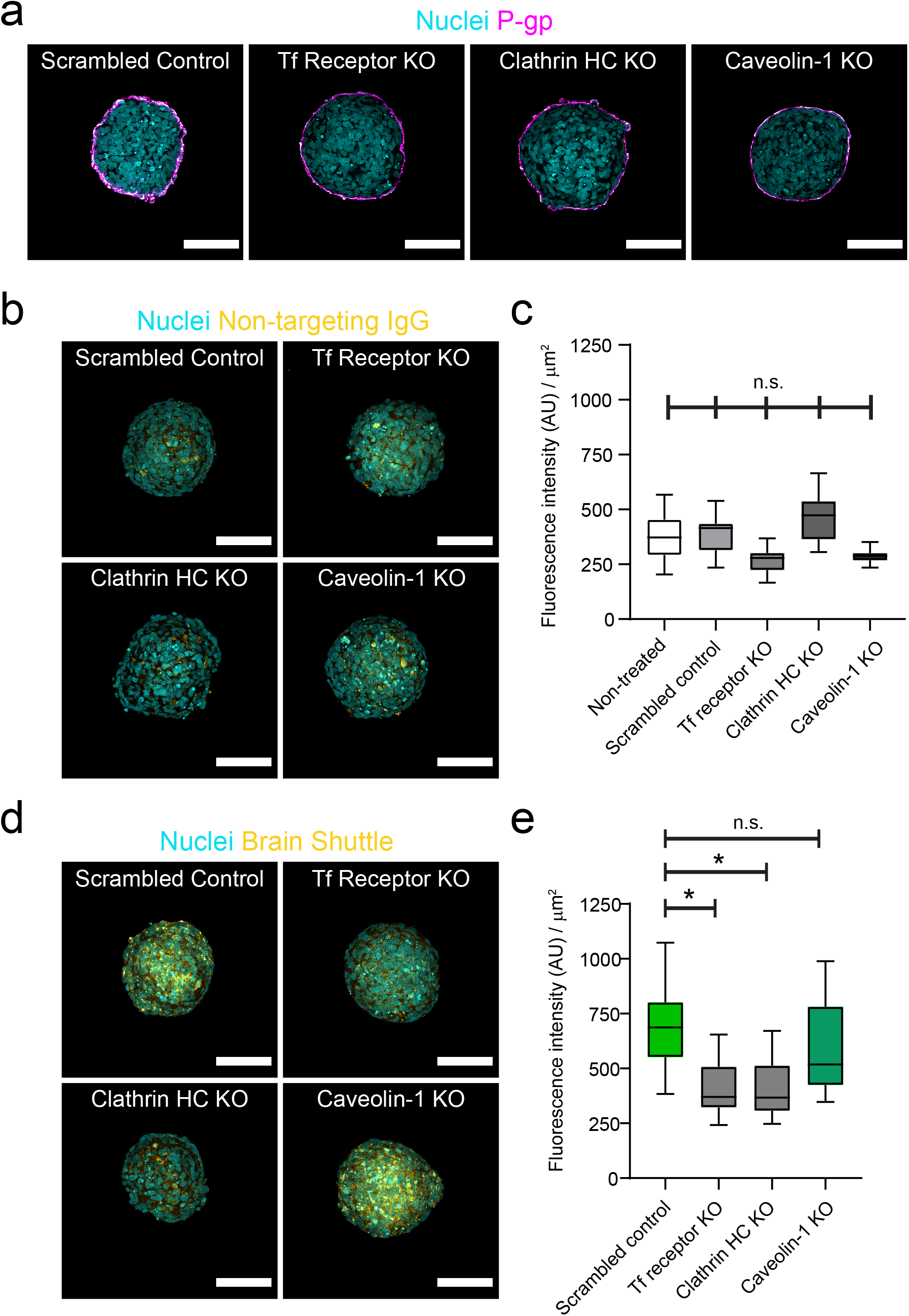
CRISPR/Cas9 gene editing in blood-brain barrier organoids to investigate the mechanisms of transcytosis. **a,** Representative confocal images of blood-brain barrier organoids assembled with hCMEC/D3 Cas9/Scrambled gRNA control or knockout cells. Brain endothelial cells are labelled with an anti-P-gp antibody (magenta) and are homogeneously distributed at the surface and forming a continuous monolayer at the periphery of the organoid core. Nuclei are labelled with DAPI (cyan). Scale bar, 100 μm. **b,** Representative confocal images of blood-brain barrier organoids incubated with a non-targeting human IgG for 4 hours. Images show an overlay of human IgG signal (yellow) and nuclei labelled with DAPI (cyan). Scale bar, 100 μm. **c,** Quantification of IgG intensity within blood-brain barrier organoids. Graph shows boxplots with interquartile ranges and median. Lines show the 5th and 95th percentiles. Differences between treatments were not statistically significant (p = 0.344) as evaluated by one-way ANOVA of 30 organoids per condition in n = 2 independent experiments. **d,** Representative confocal images of blood-brain barrier organoids assembled with hCMEC/D3 Cas9 knockout cells incubated with a human Brain Shuttle for 4 hours. Images show an overlay of human IgG signal (yellow) and nuclei labelled with DAPI (cyan). Scale bar, 100 μm. **e,** Quantification of IgG intensity within blood-brain barrier organoids. Graph shows boxplots with interquartile ranges and median. Lines show the 5th and 95th percentiles. Differences between the scrambled control and transferrin receptor or clathrin heavy-chain knockout organoids were statistically significant (*, p < 0.05) whereas the difference between the scrambled control and caveolin-1 knockout organoids was not statistically significant (p = 0.609). Comparisons were evaluated by one-way ANOVA followed by Dunnett’s test for multiple comparisons of ~ 50 organoids per condition in n = 3 independent experiments.

## Discussion

A major obstacle for accelerating the development of new brain targeting biologics is the lack of high-throughput *in vitro* BBB models that are translatable to humans. Here, we bridged this gap by using a novel bioengineering approach relying on hydrogel-based patterned microcavity arrays to grow homogeneous organoids [28,37,38]. Previous work established that BBB organoids recapitulate key properties of the BBB observed *in vivo* [25–27] and constituted a major advance in the field. The model and workflow we developed confirms the key properties and functionality of BBB organoids and builds upon previous work by making substantial improvements to facilitate its widespread adoption for drug discovery. First, microwell arrays increased the BBB organoid yield more than 50 fold per experiment compared to previous protocols using agarose [26,27] or hanging-drop culture [24,25]. This yield increase is accompanied by a substantial reduction in experimental handling time, as hydrogel production can be automated [28] and replaces manual preparation of individual plates [27]. Second, BBB organoid formation in arrays was highly reproducible between independent experiments, whereas assembly in agarose resulted in inter-experimental variation of up to 40% in organoid size. Third, we used an automated image analysis script on confocal microscopy images to measure the accumulation of antibodies within organoids. This streamlined workflow enabled the measurement of nearly 10 times more organoids compared to previous studies [25,27]. This larger data set allowed us to robustly estimate the dynamics of human Brain Shuttle transcytosis. Interestingly, accumulation of human Brain Shuttle in BBB organoids reached a steady-state around 60 minutes (Figure 3d), which could be the result of receptor saturation or the balance between transcytosis and efflux/recycling [39,40]. Via this workflow, various scenarios can be evaluated in more detailed mechanistic studies with the same *in vitro* platform. Fourth, the transport assay in BBB organoid arrays allows the characterization of native, non-labeled antibodies, whereas previous work measured BBB-transport of biologics using molecules that were covalently coupled with organic fluorophores [18,26]. While detection of fluorescent conjugates with microscopy requires no additional sample preparation, it does require the prior chemical labelling of each one of the molecules in the test set, which may not be feasible during lead identification and optimization phases. On the other hand, a limitation of detecting non-labelled antibodies by immunofluorescent labeling as done in this study is that it prevents absolute quantification of the amount of transcytosed molecules. Altogether, we consider these improvements make BBB organoid arrays an optimal platform for drug discovery to identify biologics that can cross the BBB in a scalable manner. The use of high throughput human *in vitro* models of the BBB will advance translatability of early discovery work into the clinic.

The reproducibility and high sensitivity of the transport assay in BBB organoid arrays make them a useful tool to dissect the molecular mechanisms of transcytosis. We showed that CRISPR-based gene editing can be combined with BBB organoid arrays to evaluate the role of individual genes in receptor-mediated transcytosis. We investigated caveolin-1 depletion as it has been proposed to be a key regulator of transcytosis across the BBB [20,21,41]. However, our data shows that clathrin, but not caveolin-1, is required for transferrin-receptor mediated transcytosis of a human Brain Shuttle. This finding supports the presence of multiple transcytosis pathways with different molecular machineries in brain endothelial cells [42]. Overall, we consider this approach will be useful to evaluate whether different formats (e.g. bispecific antibodies, single domain antibodies, multivalent coated particles) or receptors utilize the same transport pathways across the BBB.

## Conclusions

Human BBB organoid arrays are a high-throughput platform to generate homogeneous and reproducible BBB organoids that allow the identification and characterization of antibodies undergoing receptor-mediated transcytosis. We show here that combining high-throughput organoid analysis and gene editing technologies can lead to novel insights into the molecular regulation of transcytosis across the BBB. Implementation of transport assays with BBB organoid arrays will help accelerate the discovery and development of new brain targeting modalities.

## List of abbreviations

ALPL: alkaline phosphatase
BBB: blood-brain barrier
BEC: brain endothelial cell
CRISPR: clustered regularly interspaced short palindromic repeats
CNS: central nervous system
EDTA: ethylenediaminetetraacetic acid
FBS: fetal bovine serum
FITC: fluorescein isothiocyanate
GA-1000: gentamicin sulfate-amphotericin
GFAP: glial fibrillary acidic protein
HA: human astrocytes
HBVP: human brain vascular pericytes
hCMEC/D3: human cerebral microvascular endothelial D3 cells
hEGF: human epidermal growth factor
hFGF-B: human fibroblast growth factor B
IF: immunofluorescence
NG2: neural/glial antigen 2
Mfsd2a: major facilitator superfamily domain-containing protein 2
PBS: phosphate buffered saline
PEG: polyethylene glycol
PGP: p-glycoprotein 1
R3-IGF-I: recombinant insulin-like growth factor-I
TfR: transferrin receptor
VEGF: vascular endothelial growth factor
WB: western blot

## Declarations

### Ethics approval

Not applicable

### Consent for publication

Not applicable

### Availability of data

The datasets generated and/or analyzed during the current study are available from the corresponding author upon reasonable request.

### Competing interests

C.S., M.D., A.G., E.L., H.K., J.N. and R.V. were employed by Roche during the execution of this study. N.B. and S.H. are named as inventors on patents of the hydrogel technology used in this study. S.H., N.B. are shareholders in SUN bioscience SA, which is commercializing those patents.

### Funding

C.S. was supported by a Roche Postdoctoral Fellowship (2018-2020, RPF-ID:491).

### Authors’ contributions

S.H., N.B., and C.C. designed and fabricated hydrogel microwell plates and optimized them for BBB organoid growth. C.S. and M.D. performed experiments with BBB organoid arrays. A.G. developed the automated imaging script. C.S. developed transcytosis assay. C.S. and E.L. generated and characterized all CRISPR-Cas9 cell lines. H.K., and J.N. generated and provided critical reagents. R.V. conceived the study. C.S. and R.V. wrote the manuscript. All authors read and approved the final manuscript.

## Acknowledgements

We thank Andreas Thommen and Enrique Gómez Alcaide (pRED, PS BiOmics) for support with cell sorting of Cas9 cell lines. We thank Mohammed Ullah for guidance and support during this project.

